# Changes in urinary proteome of rats before and after noise exposure

**DOI:** 10.1101/2024.12.09.627512

**Authors:** Minhui Yang, Chenyang Zhao, Youhe Gao

## Abstract

This study established an acute noise exposure model in which 10 rats were exposed to a 119 dB noise environment for 9 hours daily over a period of 7 consecutive days. Proteomic analysis was performed on urine samples collected from the rats. A comparison of protein expression profiles before and after noise exposure identified 219 differentially expressed proteins, 30 of which are known to be associated with hearing loss. Notably, Gelsolin plays a critical role in regulating the growth and stability of the mechanosensory hair bundles in mammalian cochlear outer hair cells. Deficiency of Lysosome-associated membrane glycoprotein 1 has been reported to cause vacuolization and structural alterations in lysosomal membrane proteins in cochlear marginal cells, leading to hearing loss in mice. Similarly, SLIT and NTRK-like family member 6 deficiency is linked to delayed synaptogenesis and auditory dysfunction in mice. These findings indicate that high-decibel noise exposure induces significant changes in urinary protein expression, many of which are associated with auditory damage as previously reported.

## Introduction

A recent study released by the World Health Organization (WHO) shows that up to 1 billion young people aged 12 to 35 worldwide are at serious risk of hearing loss, mainly due to their prolonged and excessive exposure to high volume music and other entertainment sounds. With the acceleration of urbanization and industrialization in our country, the problem of noise pollution has become increasingly prominent, especially in noise intensive areas such as military factories, airports, and entertainment venues. Both workers and visitors are at great risk of hearing loss, and noise induced hearing loss has become a major cause of hearing disability that cannot be ignored.

In the diagnosis of hearing loss, temporary threshold shift (TTS) and permanent threshold shift (PTS) are two common conditions. Although histological methods are considered the gold standard for diagnosis through quantitative counting of synaptic markers, their complexity and invasiveness make it difficult to widely apply in clinical practice.

Traditionally, research on hearing loss has relied on methods such as tissue sectioning, extracting total RNA and total protein from ear tissues or cells. However, an unprecedented research field - urinary proteomics, is gradually emerging. This field has shown extraordinary potential in the discovery and analysis of biomarkers, especially in the field of hearing loss, which has been almost unexplored before. The unique advantage of urine as a biomarker source is that it is not strictly bound by physiological homeostasis mechanisms, allowing it to more sensitively capture subtle biochemical fluctuations in the body. More importantly, the urine collection process is non-invasive and convenient, greatly improving its practicality and acceptability in biomarker research.

Currently, numerous scientific studies have confirmed that proteins in urine can serve as biomarkers for various neurological disorders in the brain. This study is based on this background and aims to explore the mechanism of noise induced hearing impairment from the perspective of urine proteomics. This innovative research path is not only expected to reveal new mechanisms of noise induced hearing impairment, but also provides an unprecedented opportunity to develop new therapies for this type of hearing impairment.

## 2. Materials and Methods

### 2.1 Experimental animals and model establishment

Twenty SPF grade male Wistar rats (170-190 g) were purchased from Beijing Weitonglihua Experimental Animal Technology Co., Ltd. All animals were housed in a standard environment (room temperature 22 ± 1 □, humidity 65% -70%). The animal experiments were reviewed and approved by the Ethics Committee of the School of Life Sciences, Beijing Normal University, with the code CLS-EAW-2020-034. Method for model establishment: 20 mice were randomly divided into two groups. One group of 10 rats was subjected to noise stimulation and exposed to 119 decibels of noise for 9 hours for 7 days. The other 10 were raised under normal conditions.

### 2.2 Handling of urine samples

Before the experiment, the experimental group was placed in a rat metabolic cage for overnight urine collection. After noise stimulation, both the experimental group and the control group were placed in the rat metabolic cage for overnight urine collection.

When processing urine samples, first centrifuge them at 4 □ with a centrifugal force of 12000g for 30 minutes. Then, use three to five times the volume of ethanol to precipitate the components in each 15ml urine sample at -20 □. This process needs to be completed overnight. The next day, the protein precipitate was centrifuged again at 12000g and dissolved in lysis buffer containing 8 mol/L urea, 2 mol/L thiourea, 50 mmol/L Tris, and 25 mmol/L dithiothreitol. Finally, the protein content in the supernatant was quantitatively analyzed using the Bradford analysis method.Next, digest a total of 100 micrograms of protein using trypsin. Load the proteins in each sample into a 10 kDa filtration device, wash twice with urea buffer and 25 mmol/L NH4HCO3 solution, and then reduce with 20 mmol/L dithiothreitol at 37 □ for 1 hour. Subsequently, alkylate with 50 mmol/L iodoacetamide (IAA) under dark conditions for 45 minutes. After processing, wash the sample again with UA and NH4HCO3, and digest it overnight with trypsin (enzyme to protein ratio of 1:50) at 37 □ for approximately 14 to 16 hours. The digested peptide segments were subjected to desalination treatment using the Oasis HLB assay kit and finally dried using a freeze dryer.

For subsequent analysis, the dried peptide segments were re dissolved in a 0.1% formic acid solution and diluted to a concentration of 0.5 micrograms per milliliter. In order to construct a spectral library for data independent acquisition (DIA) analysis, mixed samples (1-2 micrograms per sample) were loaded onto an equilibrated high pH reverse phase fractionation centrifuge column. Subsequently, volatile high pH elution solutions containing 8 different concentrations of acetonitrile (5%, 7.5%, 10%, 12.5%, 15%, 17.5%, 20%, and 50% acetonitrile) were added to the column to form a step gradient for elution of peptides from 8 different gradient components. The eluted sample was dried by vacuum evaporation and resuspended in 20 microliters of 0.1% formic acid solution. Finally, 2 microliters were taken from each component for liquid chromatography data dependent acquisition tandem mass spectrometry (LC-DDA-MS/MS) analysis.

### 2.3 Data analysis

Ten components were separated by reverse phase chromatography column, and mass spectrometry data were collected using DDA (Data Dependent Acquisition) mode. Subsequently, the DDA collected results were imported into Proteome Discoverer software (version 2.1) for library searching operations. Establish a DIA (Data Independent Acquisition) collection method based on the PD search results, and calculate the window width and quantity based on the m/z distribution density. Collect mass spectrometry data using DIA mode for a single peptide sample. Using Spectronaut X software to process and analyze mass spectrometry data, import raw files collected by DIA for each sample to search the database. The criterion for determining highly reliable proteins is that the q value of the peptide segment is less than 0.01, and the peak area of all fragment ions in the secondary peptide segment is used for protein quantification.

### 2.4 Statistical analysis

Each sample was subjected to 3 technical replicates, and the obtained data was used for statistical analysis. Differential protein screening was performed on urine proteins before and after exposure to noise. The criteria for screening differential proteins were: Fold change greater than or equal to 1.5 or less than or equal to 0.67, and a P-value of less than 0.05 in the paired t-test. In random grouping analysis, 20 samples are randomly divided into two groups, and the average number of differentially expressed proteins is calculated for all random combinations using the same screening criteria.

## 3. Results and discussion

### 3.1 Urine proteomic analysis

Perform LC-MS/MS tandem mass spectrometry analysis on 20 collected urine protein samples before and after noise treatment. A total of 1552 proteins (specific peptides ≥ 2, protein level FDR<1%) were identified and subjected to unsupervised cluster analysis. From these data, urine samples before and after noise treatment can be distinguished. Figure 1 shows the specific unsupervised clustering results of the samples.

**Figure 1:**
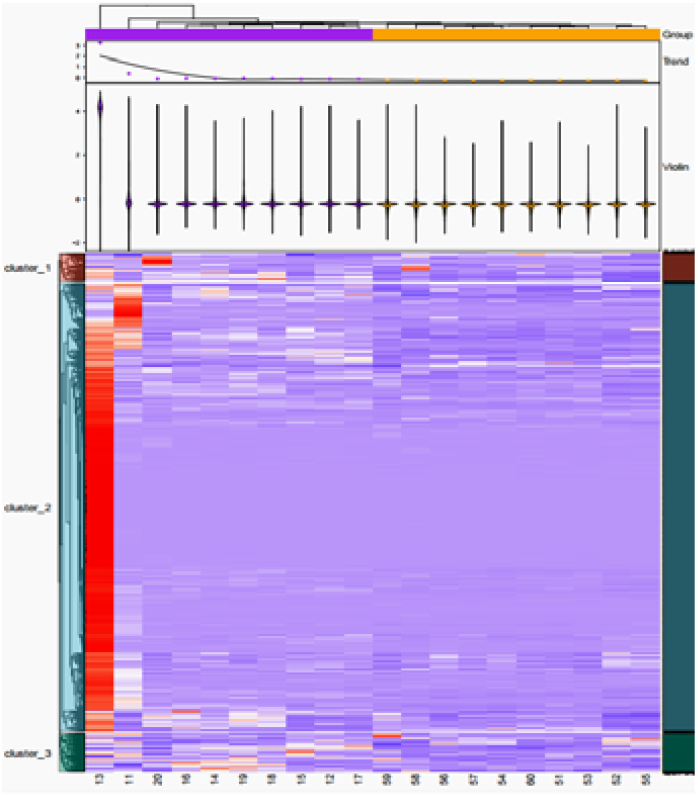
Unsupervised clustering.

### 3.2 Differential protein screening

Comparing the proteins before and after noise, a total of 219 differentially expressed proteins were identified (see Appendix), and 15 differentially expressed proteins were generated by random grouping, as shown in Table 1. The screening criteria for differential proteins are FC ≥ 1.5 or ≤ 0.67, with P<0.05. The specific information of differential proteins is listed in Table 2. By randomly matching the groups and shuffling them, an average of 15 differentially expressed proteins were obtained, proving the reliability of the differential protein results and their non random generation, as shown in Table 2.

**Table 1.**
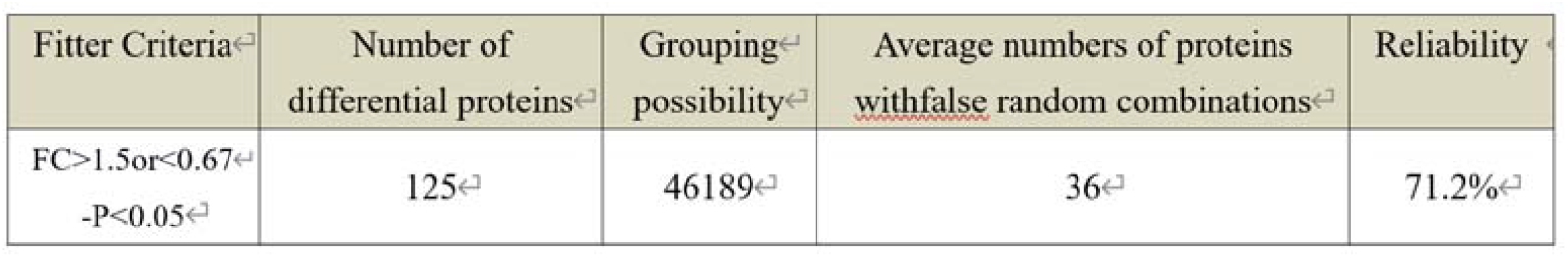
Random grouping.

**Table 2.**
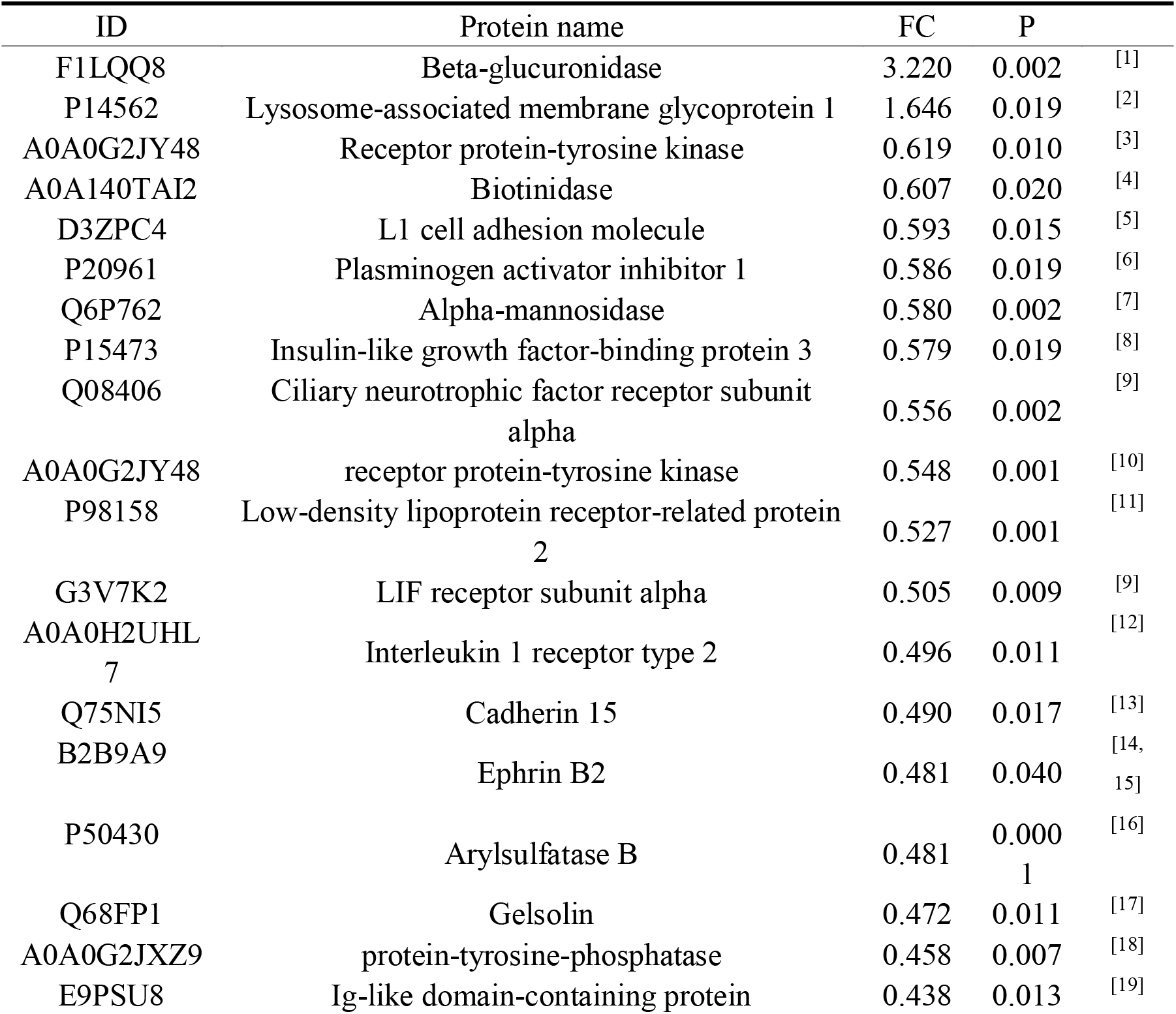
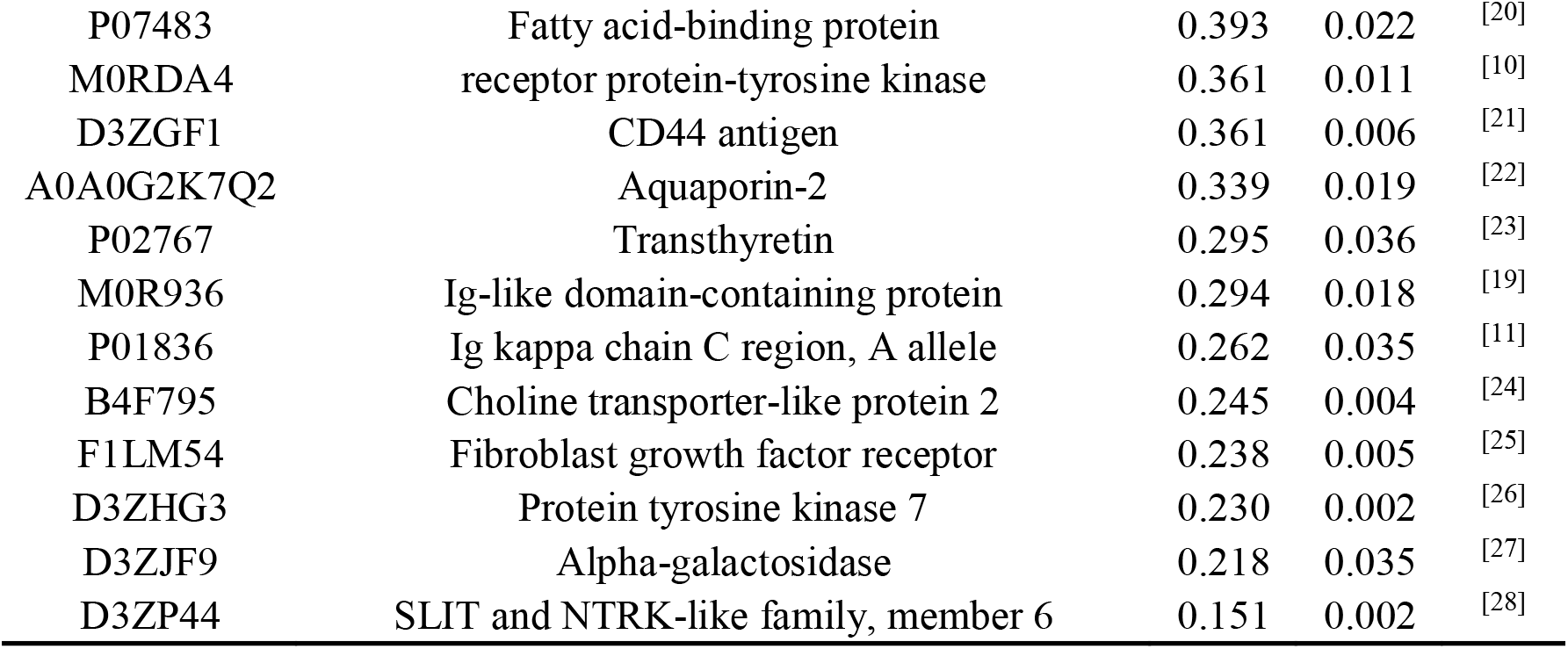
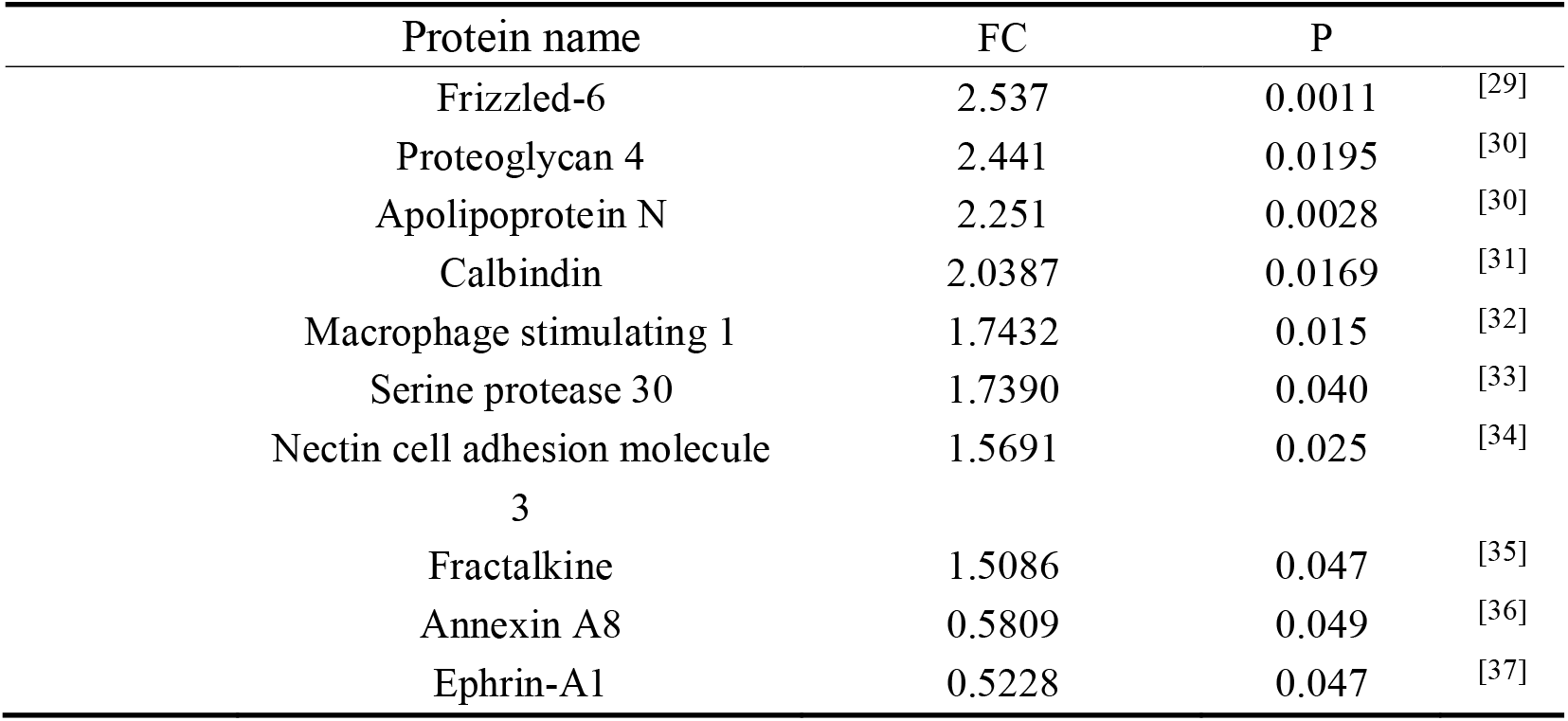
Differential Proteins.

### 3.3 Differential protein retrieval

Self comparison before and after using an open database (https://pubmed.ncbi.nlm.nih.gov/) Searching for proteins, 30 proteins related to hearing were found as shown in Table 3.

Biotinidase deficiency leads to hearing loss ^[4]^, Lysosome associated membrane glycoprotein 1 L1 cell adhesion molecules are released in the Corti organ of the mouse cochlea ^[5]^, Alpha mannosidase alpha mannosidase activity deficiency causes hearing loss ^[7]^, Insulin like growth factor binding protein 3, and lack of IGF binding protein-3 leads to hearing and learning defects [8], Low density lipoprotein receptor related protein 2 is a biomarker for vestibular schwannoma related hearing loss, while Interleukin-1 receptor type 2 is expressed in the inner ear. Overexpression of IL1R2 reduces noise induced synaptic damage and hearing loss ^[12]^. The loss of function of protein tyrosine phosphatase 1B leads to hearing loss ^[18]^, while the lack of collagen receptor DDR1 in receptor protein tyrosine kinase mice results in inner ear defects and hearing loss ^[10]^. A novel ILDR1 mutation in the Ig like domain containing protein like domain leads to high-frequency hearing loss due to disruption of the tight junctions of three cells in the inner ear ^[19]^. Protein tyrosine kinase 7 changes in tyrosine kinase B receptor levels 80 days after sound exposure and aging ^[26]^.

The study used liquid chromatography tandem mass spectrometry (LC-MS/MS) technology to search for clues of hearing loss in urine for the first time. A comprehensive and in-depth comparative analysis was conducted on the proteome of rat urine before and after exposure to noise environment. The experimental results showed that there were multiple significantly different proteins in the urine proteome of rats before and after noise exposure, many of which were identified as proteins directly related to hearing function. These proteins include but are not limited to: Beta glucuronidase, Lysosome associated membrane glycoprotein 1, Receptor protein tyrosine kinase, Biotinidase, L1 cell adhesion molecule, Plasma activator inhibitor 1, Alpha mannosidase, Insulin like growth factor binding protein 3, Ciliary neurotropic factor receptor subunit alpha, receptor protein tyrosine kinase, Low density lipoprotein receptor related protein 2, LIF receptor subunit alpha, Interleukin-1 receptor type 2, Cadherin 15, Eph rin B2, Arylsulfase B, Gelsol, protein tyrosine phosphatase, Ig like domain containing protein, Fatty acid binding protein, receptor protein tyrosine kinase, CD44 antigen, Aquaporin-2, Transthyretin, Ig like domain containing protein Ig kappa chain C region, A allele、 Choline transporter-like protein 2、 Fibroblast growth factor receptor、 Protein tyrosine kinase 7、 Alpha-galactosidase 、 SLIT and NTRK-like family, member 6_°_ Experimental data shows that hearing impairment can indeed reflect corresponding changes in the urine proteome. These differentially expressed proteins closely related to hearing function not only provide us with molecular mechanism clues for the occurrence and development of hearing impairment, but also provide potential targets for the development of effective treatment methods for hearing impairment in the future. This discovery not only deepens our understanding of the molecular mechanisms of noise induced hearing loss, but also opens up new research directions for early diagnosis and intervention of hearing impairment. Hearing loss caused by Arylsulfatase B mutation ^[16]^. Gelsolin plays a complementary role in regulating the growth and stability of mechanosensory hair bundles in mammalian cochlear outer hair cells ^[17]^. FABP7 deficiency can alleviate cochlear damage after noise exposure ^[20]^. The deletion of DDR1 gene in mice is associated with severe decline in auditory function and significant structural changes in the inner ear ^[10]^. CD44 is expressed in the inner ear epithelium and is directly related to hearing ^[21]^. Aquaporin 2 shows early and specific expression patterns in the developing mouse inner ear, indicating that this aquaporin plays an important role in the development of hearing ^[22]^. Transthyroid hormone protein amyloidosis accompanied by headache, hearing loss, and peripheral neuropathy ^[23]^. LC44A2 (solute carrier 44a2), also known as CTL2 (choline transporter like protein 2), is expressed in many supporting cell types of the cochlea and is associated with hair cell survival and antibody induced hearing loss [24]. Synaptic delay occurs in Slitrk6 deficient animals, and mutant mice exhibit auditory function deficits that reflect human phenotypes ^[28]^.

Compared with 10 rats that did not listen to noise, a total of 46 differential proteins were identified, but 49 differential proteins were generated by random grouping, with no significant difference in number. Possible reasons could be the influence of individual differences or the impact of living environment on rats themselves. But proteins directly related to hearing loss were still found among the 46 differentially expressed proteins. The reason may be the influence of individual differences or the impact of living environment on rats.

Frizzled 6 regulates the Wnt signaling pathway in mice, which in turn regulates cochlear hair cell regeneration ^[29]^. The absence of Proteoglycan 4 leads to impaired spatial coupling between pre - and postsynaptic elements, resulting in hearing loss ^[30]^. There is a direct correlation between apolipoprotein and sensorineural hearing loss ^[38]^. Calcium binding proteins regulate the concentration of calcium ions in neurons during auditory conduction. The gene mutation of alpha-1 type XI collagen body causes type 2 Stickler syndrome, which has been found in the vitreous, cartilage, and cochlea of mice. The characteristics of this disease are typical eye abnormalities and auditory dysfunction ^[39]^. Cochlear macrophages regulate cochlear inflammation and may have the potential to protect hearing function from damage, including acoustic overstimulation. The number of cochlear macrophages increases 3-7 days after acoustic stimulation ^[32]^. In the auditory epithelium of the cochlea, sensory hair and supporting cells are arranged in a checkerboard mosaic pattern, which is conserved across a wide range of species. Cell adhesion molecules nectin-1 and nectin-3 are necessary for the formation of this pattern. The checkerboard pattern is considered necessary for auditory function, but has never been examined. Here, we demonstrate the importance of the checkerboard cell pattern in the survival and function of sensory hair cells in the cochlear auditory epithelium of nectin-3 knockout (KO) mice. Nectin-3 KO mice exhibited progressive hearing loss associated with degeneration of hair cells attached abnormally through apoptosis. Apoptotic hair cell death is caused by the disorder of tight junctions between hair cells. Our research suggests that the checkerboard shaped cell pattern in the auditory epithelium provides the structural basis for ensuring the survival of cochlear hair cells and hearing function ^[34]^. Fractalkine has a regulatory effect in the nervous system, which may affect the function or protection of auditory neurons ^[35]^. The use of membrane-bound protein A1 derived peptides for drug therapy can prevent cisplatin induced hearing loss ^[36]^. EphA2 regulates its localization in the inner ear by interacting with pendrin, affecting hearing function. Its mutations or functional impairments may lead to hearing loss in patients with Pendred syndrome and vestibular aqueduct enlargement (EVA) ^[37]^.

## 4. Conclusion

This study used proteomics technology for the first time to investigate the effects of noise exposure on rats in urine proteomics. The experimental results showed that there were multiple significantly different proteins in the urine proteome of rats before and after noise exposure, many of which had been reported to be directly related to hearing function. The urinary proteome is a new window for studying hearing.

## Appendix 1

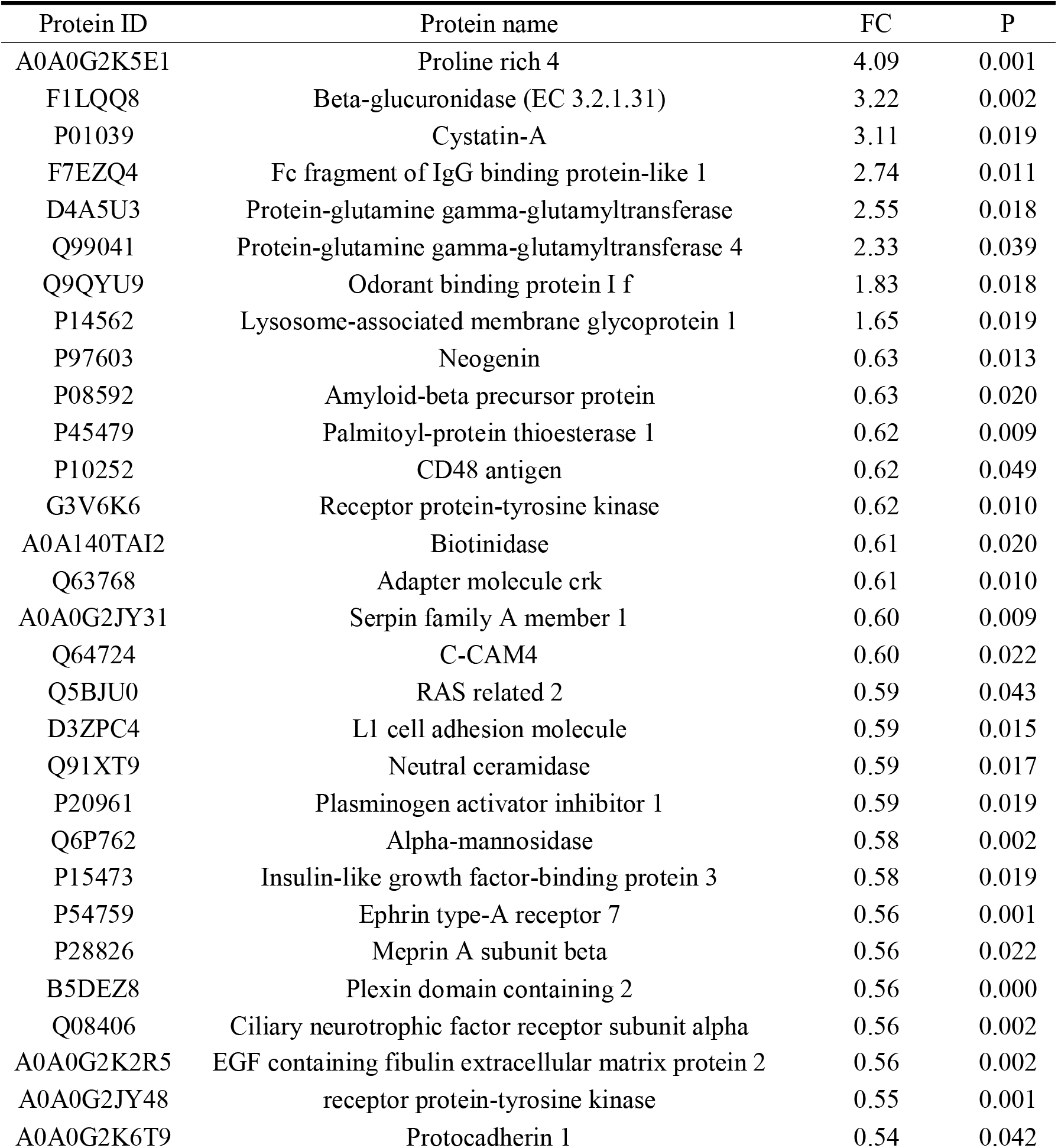
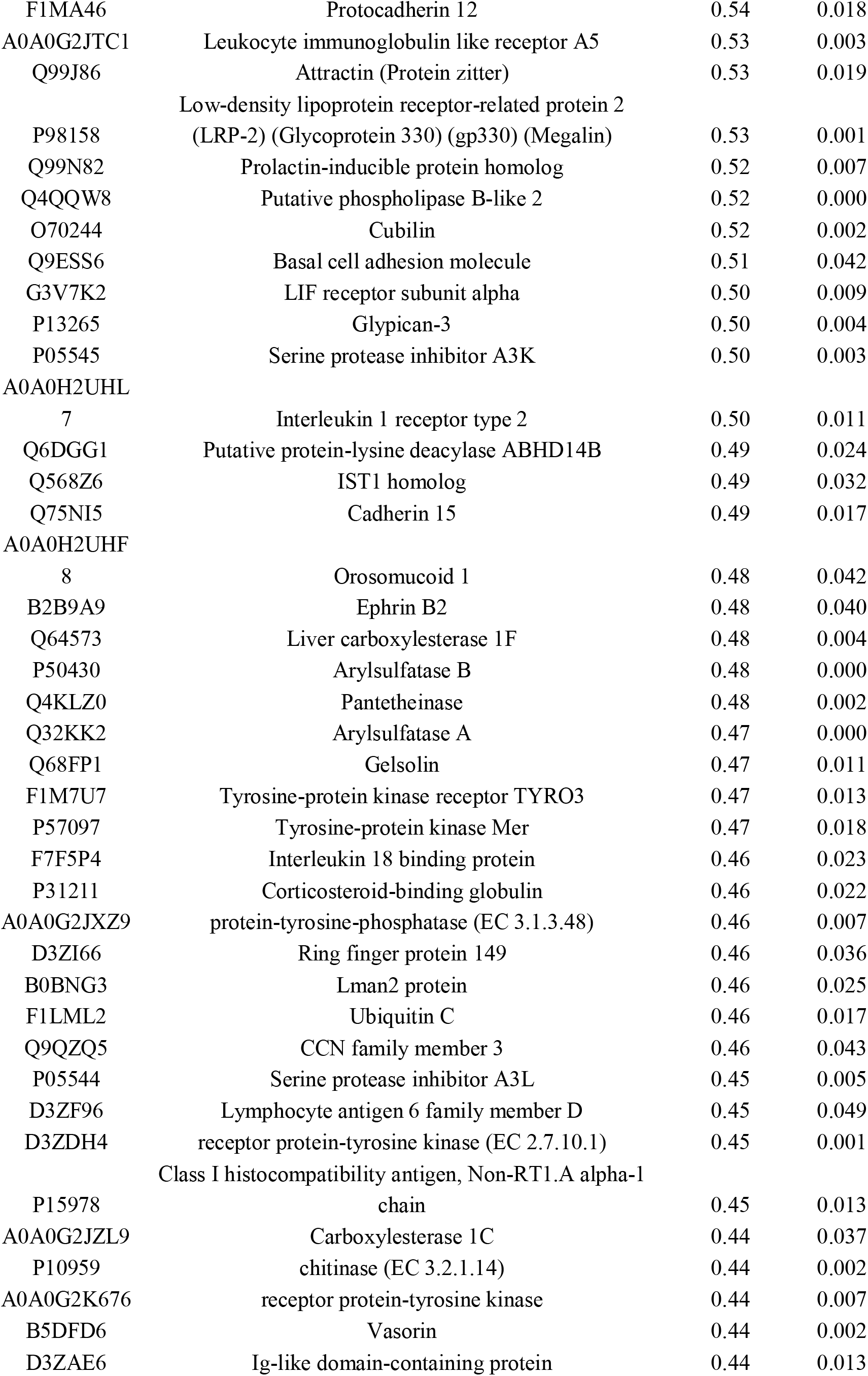
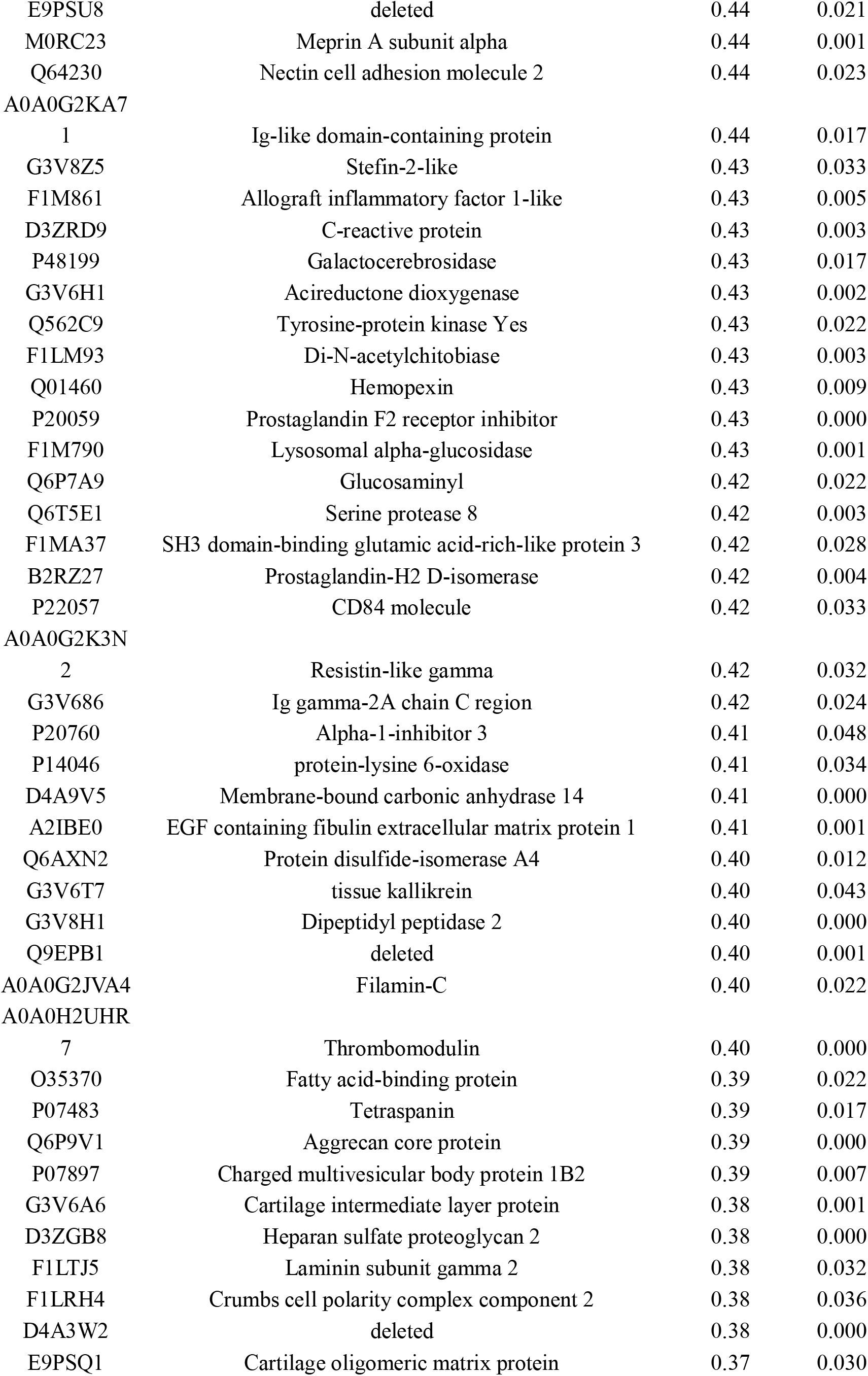
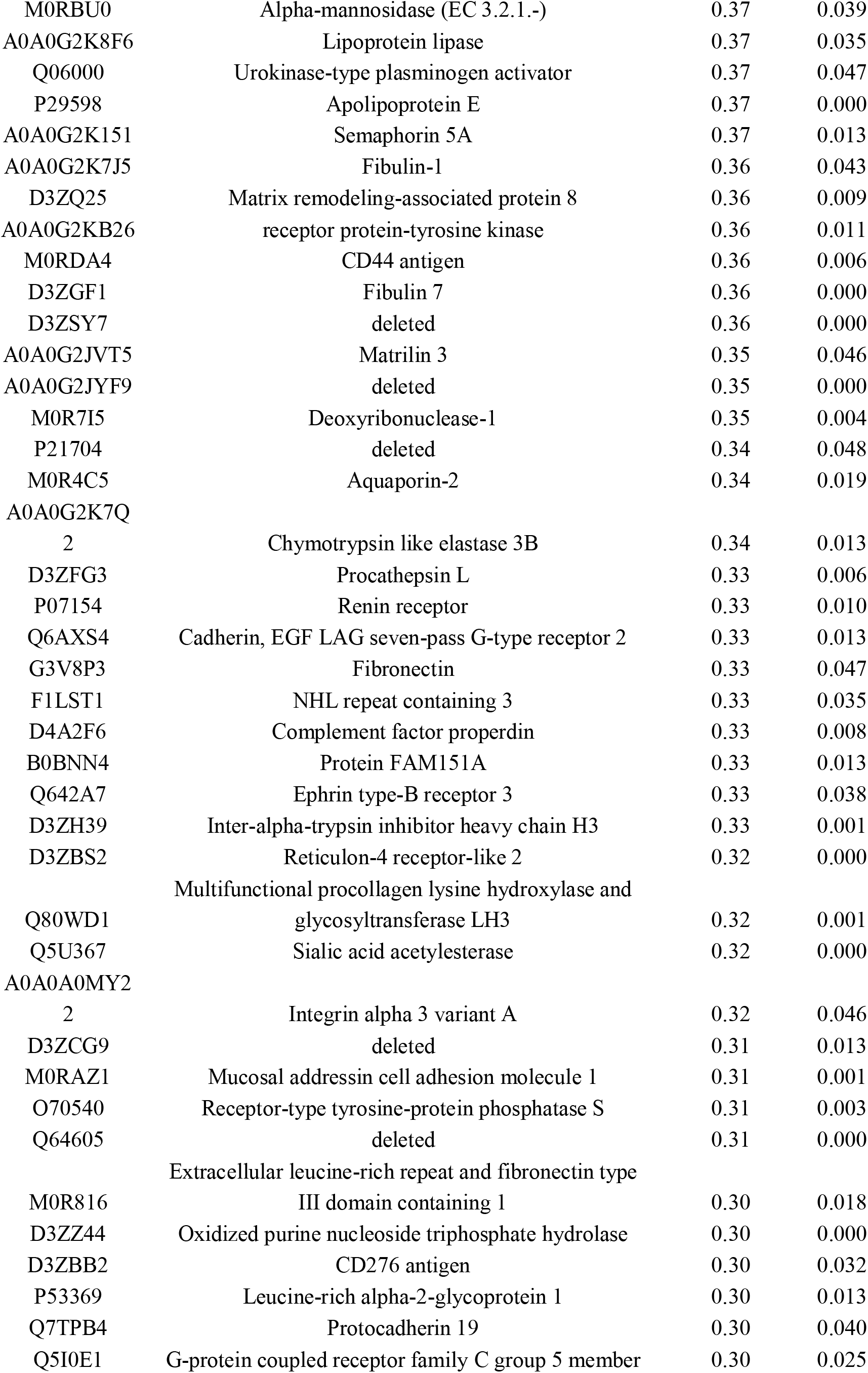
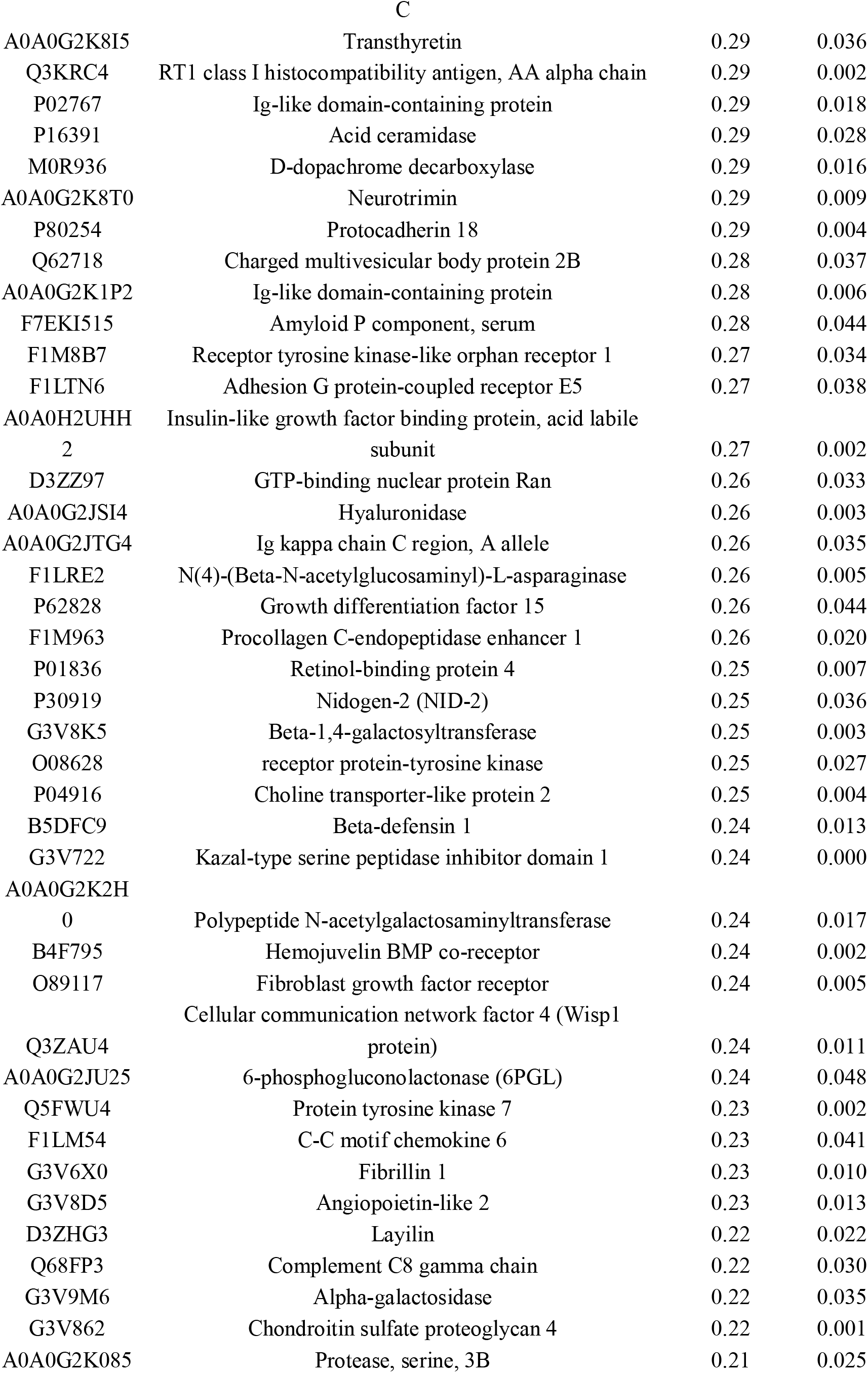
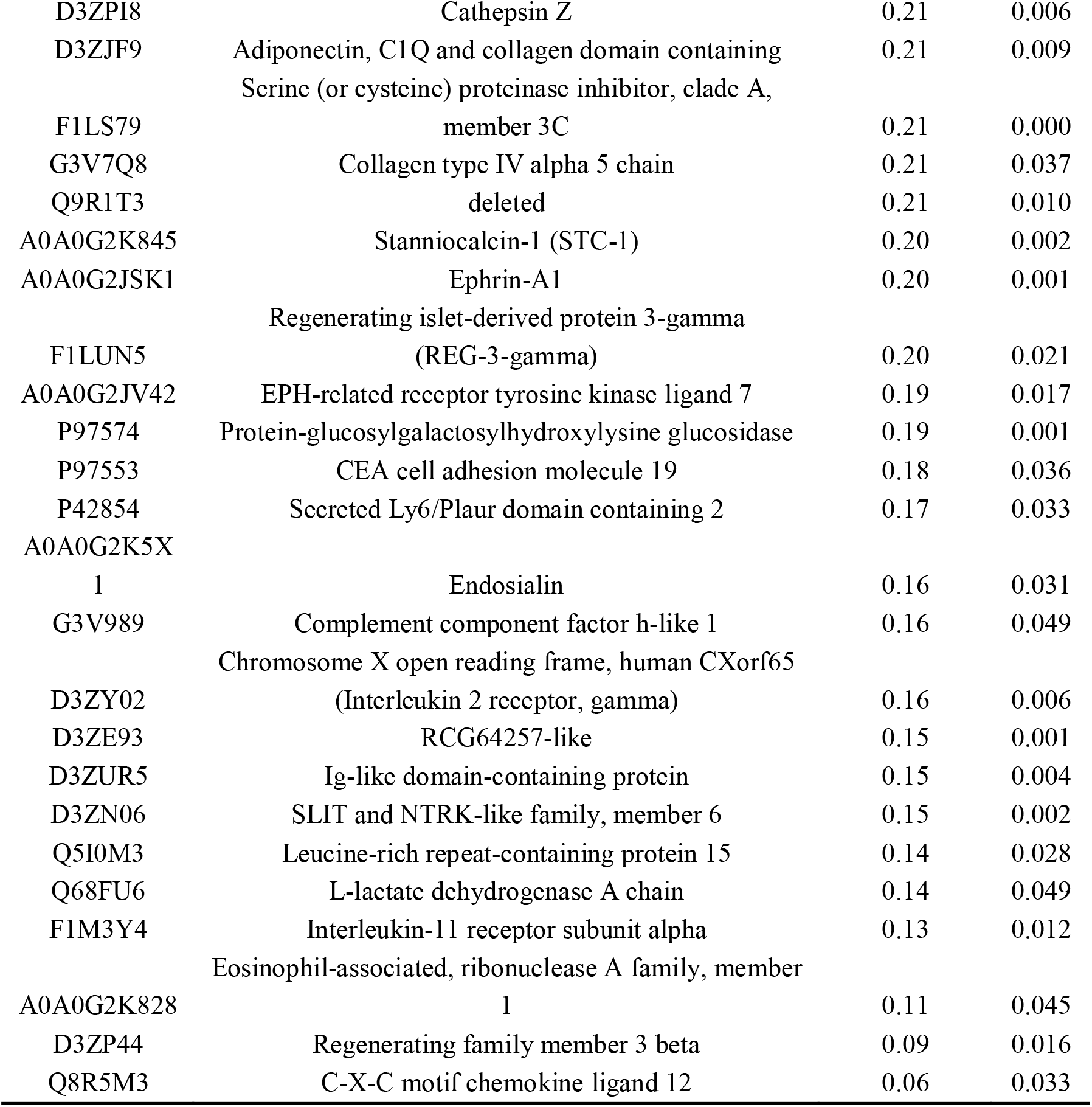

